# Cross-correlation increases sampling in diffusion-based super-resolution optical fluctuation imaging

**DOI:** 10.1101/2024.04.01.587586

**Authors:** Jeanpun Antarasen, Benjamin Wellnitz, Stephanie N. Kramer, Surajit Chatterjee, Lydia Kisley

**Author notes:** The authors contributed equally to this work (Jeanpun Antarasen, Benjamin Wellnitz).

## Abstract

Correlation signal processing of optical three-dimensional (x, y, t) data can produce super-resolution images. The second order cross-correlation function *XC*_2_ has been documented to produce super-resolution imaging with static and blinking emitters but not for diffusing emitters. Here, we both analytically and numerically demonstrate cross-correlation analysis for diffusing particles. We then expand our fluorescence correlation spectroscopy super-resolution optical fluctuation imaging (fcsSOFI) analysis to use cross-correlation as a post-processing computational technique to extract both dynamic and structural information of particle diffusion in nanoscale structures simultaneously. We further show how this method increases sampling rates and reduces aliasing for spatial information in both simulated and experimental data. Our work demonstrates how fcsSOFI with cross-correlation can be a powerful signal-processing tool to resolve the nanoscale dynamics and structure in samples relevant to biological and soft materials.

**TOC Graphic:** 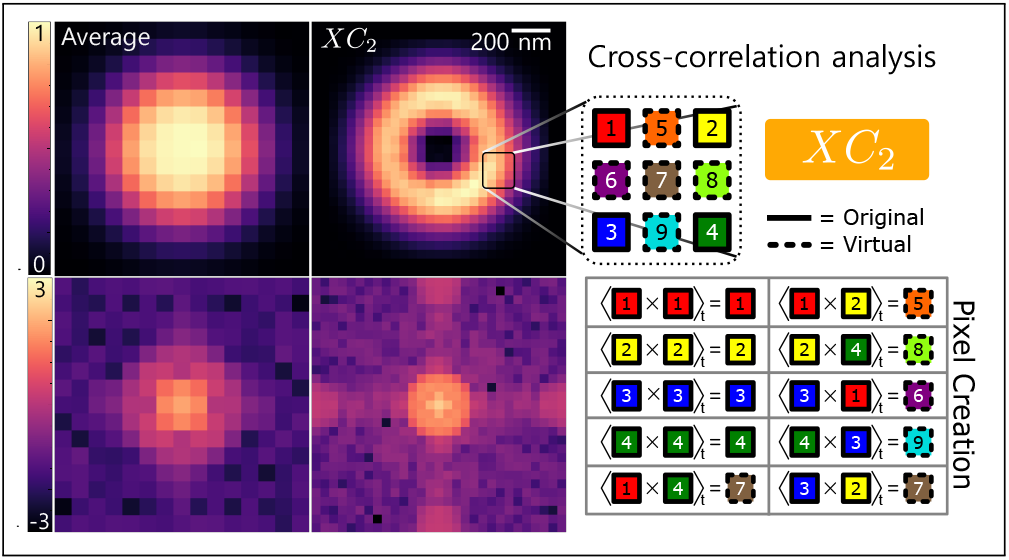

## Introduction

Correlation analysis of fluctuations measured by fluorescence microscopy is a powerful technique to determine the dynamics of biomolecules and resolve structural features at the nanoscale. ^1–6^ Mathematically correlating the intensity fluctuations in fluorescence signals over time through a single-point measurement with confocal microscopy can obtain dynamics with fluorescence correlation spectroscopy (FCS).^7–9^ Imaging FCS is a variation of FCS that incorporates either a sensitive electron-multiplying charge-coupled device (EMCCD)^3,10–12^ or a scientific complementary metal–oxide–semiconductor (sCMOS) ^12^to obtain the diffusion of biomolecules at each pixel within a two-dimensional camera array. Super-resolution optical fluctuation imaging (SOFI) is another powerful method using a wide-field camera that evaluates fluorescence fluctuations at each pixel using auto- and cross-correlation functions to spatially resolve a sub-diffraction-limited image of biomolecule structure.^13–19^ SOFI has been proven to experimentally work with static emitters such as quantum dots,^13,20^organic dyes,^15^ and fluorescent proteins^21^ detected with an EMCCD or sCMOS camera.^22^ Correlation is the mathematical backbone of FCS and SOFI, which allows for either an understanding of the diffusion dynamics or the super-resolved structure of biomolecules and biomaterials, respectively. Imaging FCS and SOFI use the size of the camera’s pixel to spatially resolve physical information, while in FCS the spatial resolution is determined by the focal volume. The distinction between imaging FCS and SOFI is only the source of fluctuation: diffusion or blinking.

The hybrid method fcsSOFI has shown that auto-correlation analysis with diffusing emitters can achieve super-resolution images similar to conventional SOFI which uses static blinking emitters.^23^ fcsSOFI precisely determines both where and how fast molecules are diffusing rather than one or the other. By measuring the fluorescent signals at each pixel over a set timescale (Figure 1a), we can use the intensity fluctuations to calculate the auto-correlation over time at each pixel (Figure 1b, pixels with solid outlines). We can then extract diffusion coefficients at each position with imaging FCS analysis (Figure 1c).^23^ Constructing an fcsSOFI image is done by using the SOFI spatial information as a grayscale, saturation transparency map on top of the colored diffusion map generated from FCS analysis (Figure 1d). We can then determine the dynamic and structural information of diffusing particles within one image and can further inspect the distribution of diffusion rates as a cumulative distribution function (CDF). Our group has previously demonstrated the use of fcsSOFI to accurately study the dynamics of fluorescent beads, polymers, and biomolecules, such as dextran and fibrinogen through experiment and simulation, and in extracellular matrix (ECM)-like materials such as collagen-mimicking nano-channels, porous agarose, and polyacrylamide hydrogels.^24,25^ Overall, fcsSOFI can produce a super-resolution image to spatially understand molecular diffusion within heterogeneous structures.

**Figure 1:**
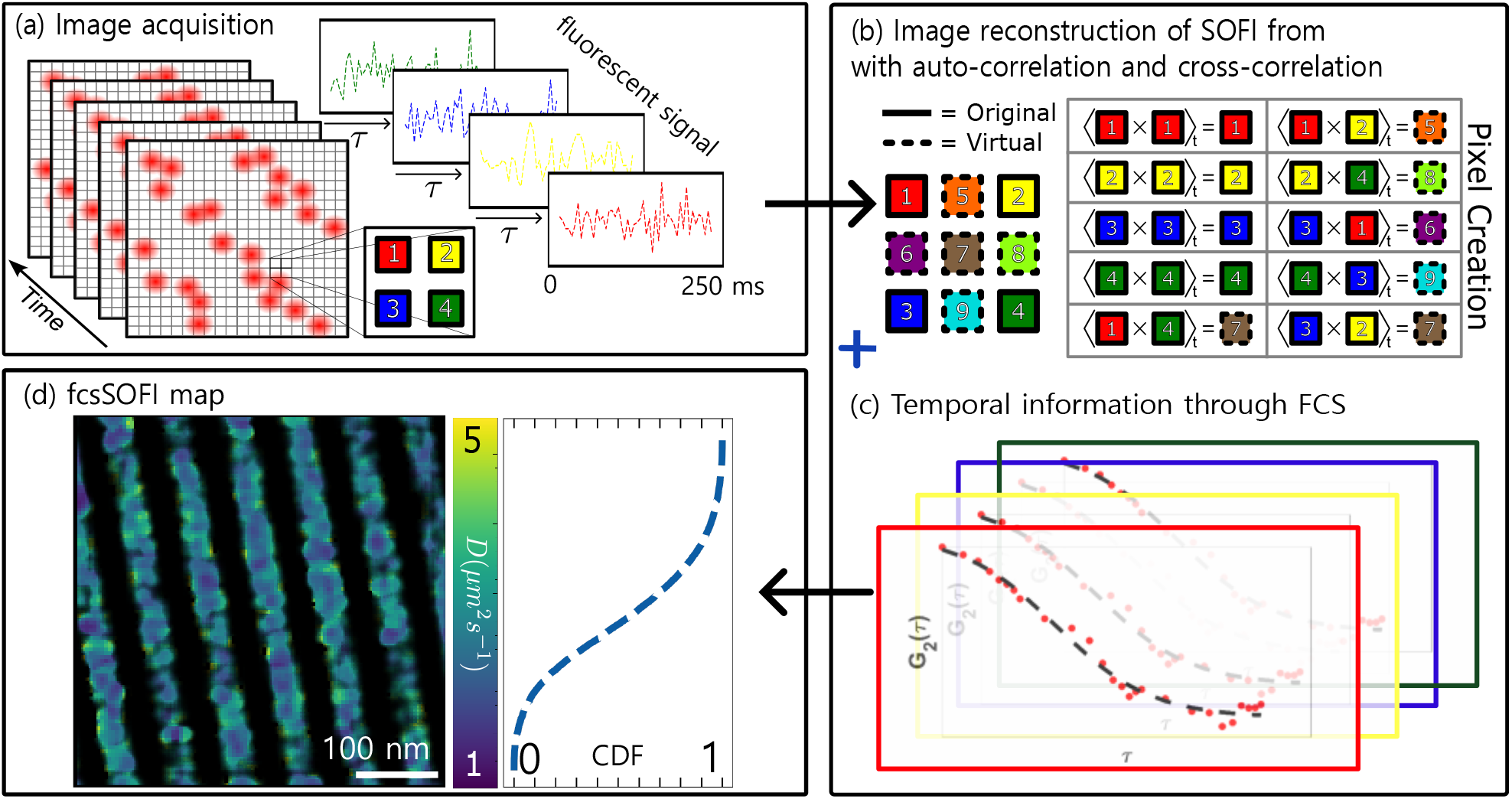
Schematic representation of fcsSOFI with cross-correlation analysis, which can simultaneously characterize the dynamics of biomolecules in confined structure. **(a)** Image acquisition to obtain the fluorescent signal at each pixel recorded on a 2D camera (e.g. EMCCD or sCMOS). **(b)** Using raw signals, cross-correlation analysis creates virtual pixels to increase pixel density. **(c)** Correlation analysis extracts diffusion information through imaging FCS. **(d)** Combination of FCS and SOFI creates a map representing the structure and the diffusion coefficient at every pixel from the fitting of auto-correlation curves. The cumulative distribution function (CDF) presents the diffusion coefficient *D* distribution for all pixels within the full-size image and quantifies the average diffusion rate in one image.

Both the fluorescent intensity and the pixel size limit the diffusion dynamics and spatial information obtainable in auto-correlation fcsSOFI. Having an analyzable fluorescent signal that can produce accurate physical information will depend on having an appropriate molecular concentration while also understanding the complexity and heterogeneity of the environment^26^ and the camera’s instrumental response function compared to the emitter diffusion.^27^ Moreover, using auto-correlation, the pixel size of the reconstructed image is restricted to the pixel size of the camera, which will not be beneficial for spatial features smaller than the super-resolved spatial resolution.^13,17^

Cross-correlation analysis has been used with SOFI to overcome the limitations of pixel size. Second-order cross-correlation SOFI analysis of static emitters was first introduced to resolve pixel-density challenges by effectively increasing the number of pixels two-fold with virtual pixels (Figure 1b, dashed outlines).^13,28^ Later, Fourier padding in SOFI was developed to improve the spatial resolution. ^17^However, cross-correlation and the Fourier method have been limited to fluorophores with fixed locations and independent photo-blinking-based stochastic fluctuations. The cross-correlation framework introduced by Vandenberg and Dedecker ^29^ expands traditional SOFI theory to include emitters that both photo-blink and diffuse. However, in their analysis, most of the SOFI information came from the photoblinking nature of the emitters with an on-time ratio of 9% and the diffusion rate was limited to 1 *µm*^2^*s*^−1^ or less. The fcsSOFI framework must be extended to understand crosscorrelation analysis of emitters that have no photo-blinking and diffuse at a wider range of rates due to the correlation of emitters from diffusion.

In this work, we report the spatial sampling enhancement to fcsSOFI by incorporating cross-correlation analysis and demonstrate the recovery of both diffusion and complex structural information using dynamics from simulations and experiments. Compared to conventional SOFI analysis which applies to the case of static blinking emitters or diffusing, we show proof that cross-correlation is also applicable to analyzing diffusing emitters at a wide range of rates. First, the theoretical framework and computational technique for reconstructing super-resolved images are provided. Next, our simulations demonstrate the regimes of temporal sampling and diffusion constants where cross-correlation SOFI is applicable with dynamic emitters. Moreover, with the benefit of higher spatial sampling, cross-correlation analysis shows the recovery of higher spatial frequency in the Fourier domain, which quantifies structure. We then produce fcsSOFI images of experimental data using cross-correlation to demonstrate our methods for a better visual and quantitative understanding of an ECM-like environment.

## Theory and Computational Method

### Revisiting the theory of auto-correlation SOFI and fcsSOFI

The theory behind SOFI for static^13,20,30^ and diffusing emitters^23^ has been previously explained in the literature. Still, we will first present a summary of the second-order autocorrelation (*AC*_2_) with diffusing emitters used for spatial analysis in fcsSOFI before expanding the theory using the second-order cross-correlation (*XC*_2_). In fcsSOFI, the spatial SOFI images are generated by auto-correlating the fluctuations in the fluorescent signal from diffusing emitters. First, we define the fluorescence signal, *F*, at a pixel position *r* and time *t* as

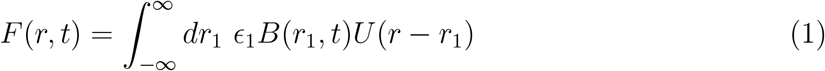

where *ϵ*_1_ and *B*_1_ represent the constant brightness of an emitter and the probability of an emitter being located at position *r*_1_, respectively.^2^ Here the point spread function (PSF) is represented as a two-dimensional Gaussian function 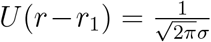 exp 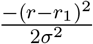 centered at an emitter’s position with standard deviation, *σ*, representing a PSF caused by the diffraction limit of light projected on a two-dimensional camera. A larger *σ* leads to a larger PSF, creating a lower-resolution image. With the model presented in Eq. 1, the optical signal fluctuates with time as particles diffuse in the image plane. ^2,23^ The signal fluctuation, *δF* can be written as

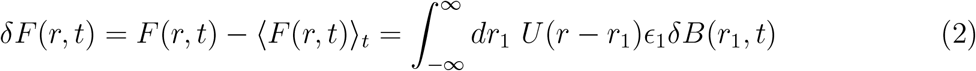

where *δB*_1_(*t*) = *B*_1_(*t*) − ⟨*B*_1_(*t*)⟩_*t*_ is the fluctuation resulting from the probability of finding emitters at *r*_1_. From Eq. 2 we can find the second-order auto-correlation in time, *AC*_2_(*r, τ*), at position *r* and exposure time (sampling rate) *τ* of Figure 2a.

**Figure 2:**
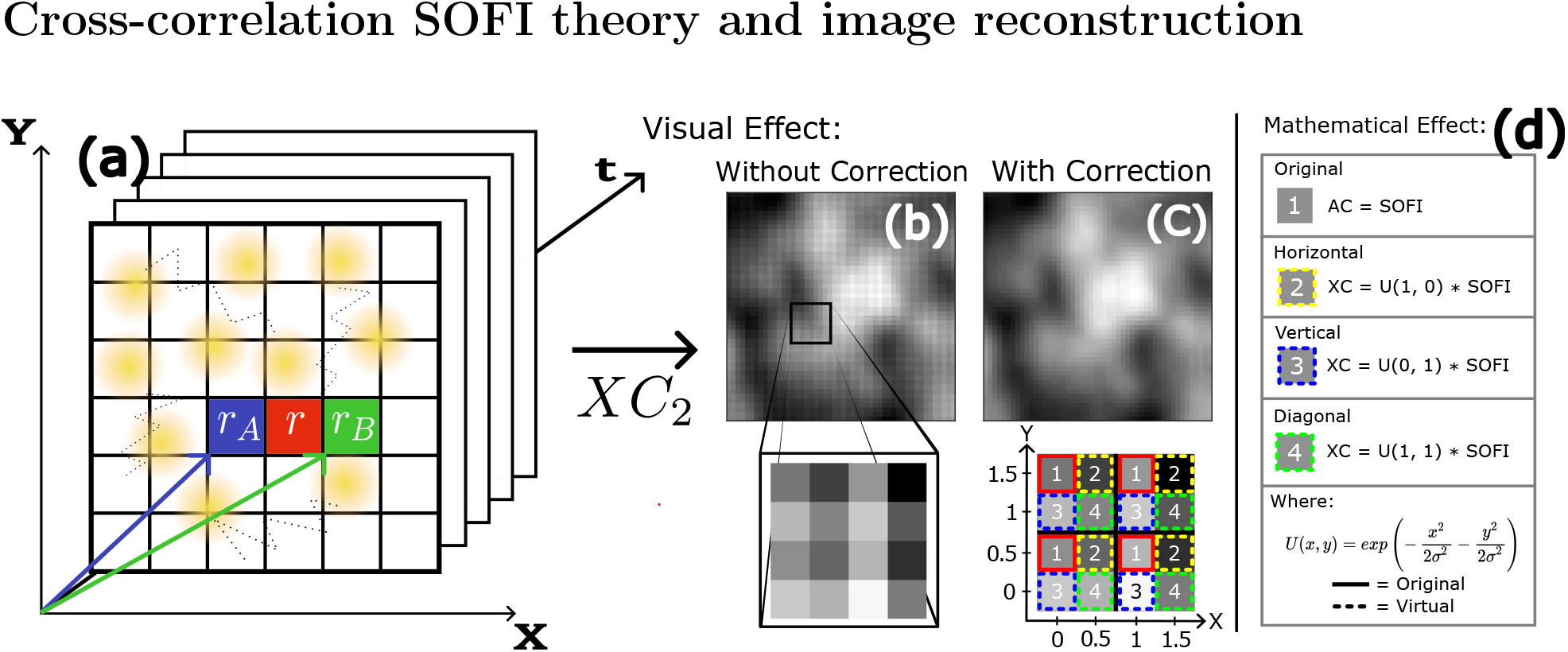
Effect of the distance factor in *XC*_2_ SOFI reconstruction at *r* from pixel position *r*_*A*_ and *r*_*B*_ on the Cartesian coordinates with a pixel size (each grid size) of 102 nm **(a)**. In **(b)**, the visual checkerboard effect can be seen on the full scale. By zooming in on a small section in **(b)**, the repetitive nature of the distance factor checkerboard effect can be found on a pixel-by-pixel level. The corrected *XC*_2_ image where the distance factor has been removed, and there is no checkerboard effect **(c)**. The mathematical representation of the distance factor where the distance between cross-correlated pixels affects SOFI values **(d)**

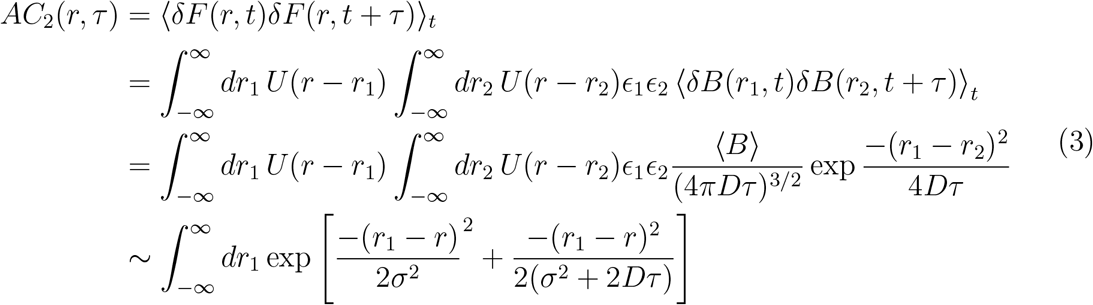

As each emitter is independent and the diffusion process is both ergodic — meaning all the accessible microstates of the locations of the particle undergoing Brownian motion are equally probable over a long period or time — and stationary — meaning the probability distribution underlying the stochastic process is invariant in time^2^ — we can analytically solve Eq. 3 by rewriting the temporal average of the fluctuation from the probability at *r*_1_ ⟨*δB*_2_(*t* + *τ*)*δB*_1_(*t*)⟩ to a spatial average that contains the diffusion information from the random-walk term 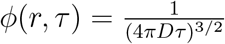 exp 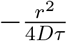, which is yielded from the diffusion equation (See Supporting Information for full derivation). SOFI reconstruction, therefore, yields a higher resolution as the new PSF in Eq. 3 in the denominator of the exponential term is smaller than the diffraction-limited PSF prior to any SOFI analysis in Eq. 1. However, the resolution improvement with diffusing emitters is limited by an increasing *Dτ*, as we will explore quantitatively in more detail in our simulation.^23^ Overall, the resulting SOFI analysis of diffusing emitters results in a super-resolution image with spatial resolution improvement due to a smaller PSF with the same number of pixels as the original image, shown in Figure 1b represented by solid outlined pixels.

### Cross-correlation SOFI theory and image reconstruction

Although *XC*_2_ SOFI with static emitters is a well-developed method,^13,20,30,31^ the analytical representation of *XC*_2_ SOFI, which explains the increased resolution and 2-fold increase in pixels, has not yet been developed for strictly dynamic, diffusing emitters. Since the mathematical representation of the fluorescent signal is different when analyzing static vs.

dynamic emitters, we must derive a new analytical expression for the *XC*_2_ SOFI analysis. With an understanding of the theory behind *AC*_2_ SOFI analysis for diffusing emitters, we now can follow similar methods to derive the analytical expression of *XC*_2_ analysis between two spatial locations at position *r*_*A*_ and *r*_*B*_ with exposure time *τ* as shown in Figure 2(a). The second-order cross-correlation, *XC*_2_(*r*_*A*_, *r*_*B*_, *τ*), is expressed as

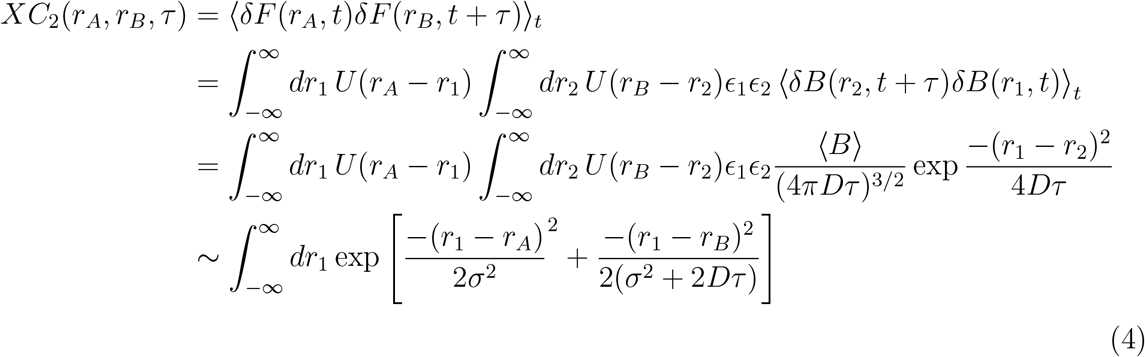

The result of Eq. 4 can be interpreted similarly to that of Eq. 3, where the new exponential term is viewed as a Gaussian PSF with a reduced *σ*. The difference between Eq. 4 and Eq. 3 is that *XC*_2_ leads to the new PSF being between positions *r*_*a*_ and *r*_*b*_. This new middle position is what gives rise to the two-fold increase in pixels for each dimension, leading to a four-fold increase in pixels overall.

When constructing a *XC*_2_ SOFI image using Eq. 4, we observe a checker-board like pattern of alternating bright and dark pixels as seen in Fig 2(b). In *XC*_2_ SOFI with static emitters, these checkerboard patterns come from a term in the *XC*_2_ equation called the distance factor.^13,30^ The distance factor can be seen in the ratio of auto-correlation 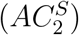 and cross-correlation 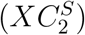 for traditional SOFI images, where *S* represents that the data is from static, photoblinking emitters.

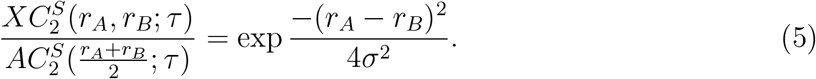

For SOFI with diffusing emitters, we follow a similar technique to determine the distance factor by analytically integrating both Eq. 3 and Eq. 4 (see Eq. S1 to S5 for a full derivation):

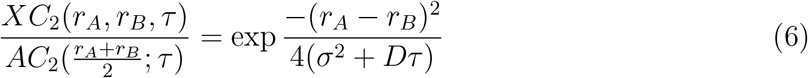

Solving the distance factor analytically is just one of many proposed methods for correcting the distance factor. The original method for correcting the distance factor in literature fit for the PSF which led to the minimum local variance, therefore smoothing any artifacts out of the image.^13^ Other proposed methods include assumptions about the variance between pixels, which however valid, lead to the artificial adjustment of pixels to fit the assumptions.^13,14^ In this work, we correct for this distance factor by adapting a method used on SOFI with static emitters which analytically solves and removes the distance factor from the image.^30^ Solving for the distance factor analytically allows us to retain the true SOFI values for future analysis and has no artificial adjustments. As seen in Figure 2d, a *XC*_2_ SOFI image consists of three different distances and, thus, three different distance factors. By calculating a *XC*_2_ SOFI image where positions *r*_*A*_ and *r*_*B*_ are chosen such that all virtual pixel positions match with real pixel positions, we obtain the distance factor at the three different distances. Then, the calculated distance factors can be used to solve for *σ*, an approximate standard deviation of the Gaussian, in Eq. 5 (the PSF in SOFI with static emitters) and 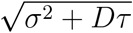 in Eq. 6. Correcting for and removing the calculated distance factors from an image reconstruction yields the inherent high-resolution SOFI information with no checkerboard pattern artifacts (Figure 2c). However, a diffusion term, *Dτ*, differentiates Eq. 6 from Eq. 5, emphasizing the significance of accounting for diffusion rate and speed of image acquisition in the calculation of distance factor. Later, we will demonstrate the regime where values of the diffusion term (*Dτ*) are applicable in *XC*_2_ SOFI reconstruction through simulations with known diffusion coefficient, *D*, and time lag, *τ* .

The *XC*_2_ of the two-pixel positions *r*_*A*_ and *r*_*B*_ yields the same numerical values as the *AC*_2_ of the pixel position at the mean position of 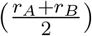 between the two *XC*_2_ pixels. After correcting images by the removal of the distance factor, *XC*_2_ can effectively be used to interpolate virtual pixels which are between the two real pixels and contain the same spatial resolution improvement as their *AC*_2_ counterparts since *AC*_2_ and *XC*_2_ differ by nothing but the distance factor. Hence, *XC*_2_ is applicable within the fcsSOFI technique in the same manner as *AC*_2_.

### Diffusion simulation method

A simulation in MATLAB is used to mimic emitter diffusion with known parameters of the diffusion coefficient, the structure the particle is diffusing within, and the PSF of the microscope that can be unknown in experimental situations where SOFI analysis would be applied.

Here, we simulate fluorescent emitters moving in three dimensions with Brownian motion and define diffusion coefficient *D* for time step *τ* with *n* number of frames. The number of particles is limited to 1-20 emitters diffusing within the 1.5 *µm* x 1.5 *µm* area of the structure to replicate the sampling in typical fcsSOFI imaging. We also set a pixel size of 0.1 *µm* x

0.1 *µm* to simulate our camera’s pixels. The randomness of the movement of each particle is generated by a random integer generator (**randi() in MATLAB**). The movement of the particles is restricted based on a binarized pore map that features accessible areas inside generated structures such as vertical channels, rings, or random pores (Figures 3a, 3e, 4a, and S2c to e respectively). If a particle hits the edge of the structure boundary, it will stop at the wall for one frame to justify confinement and randomly move in the next frame. The simulation of movement generates a ground-truth video for particles diffusing in the structures. We then apply a convolution (**conv2() in MATLAB**) of the images with a two-dimensional Gaussian function (**fspecial(‘gaussian’) in MATLAB**) to demonstrate the effect of a PSF from the diffraction limit of light in microscopy. The *σ* of the PSF is 204 nm. The brightness of a particle *ϵ*_1_ at position *r*_1_ is determined from a Poisson distribution (**poissrnd() in MATLAB**) with a constant mean to demonstrate the intensity distribution in a physical system. Therefore, our simulation includes values for *ϵ*_1_ and *U* (*r* − *r*_1_) in the analytical expression for the fluorescence signal shown in Eq. 1. Shot noise is generated by randomly pulling numbers from a Poisson distribution for each pixel position using its intensity from convolution as an average rate. Lastly, we add uniformly random read noise as background *b*(*r*) (**rand() in MATLAB**) at each pixel position *r*. During the correlation analysis, any effect of white noise is theoretically removed since the white background noise is not correlated in time. We have published the GitHub code^32^indexed on Zenodo for the settings and parameters we use to create a simulation.

**Figure 3:**
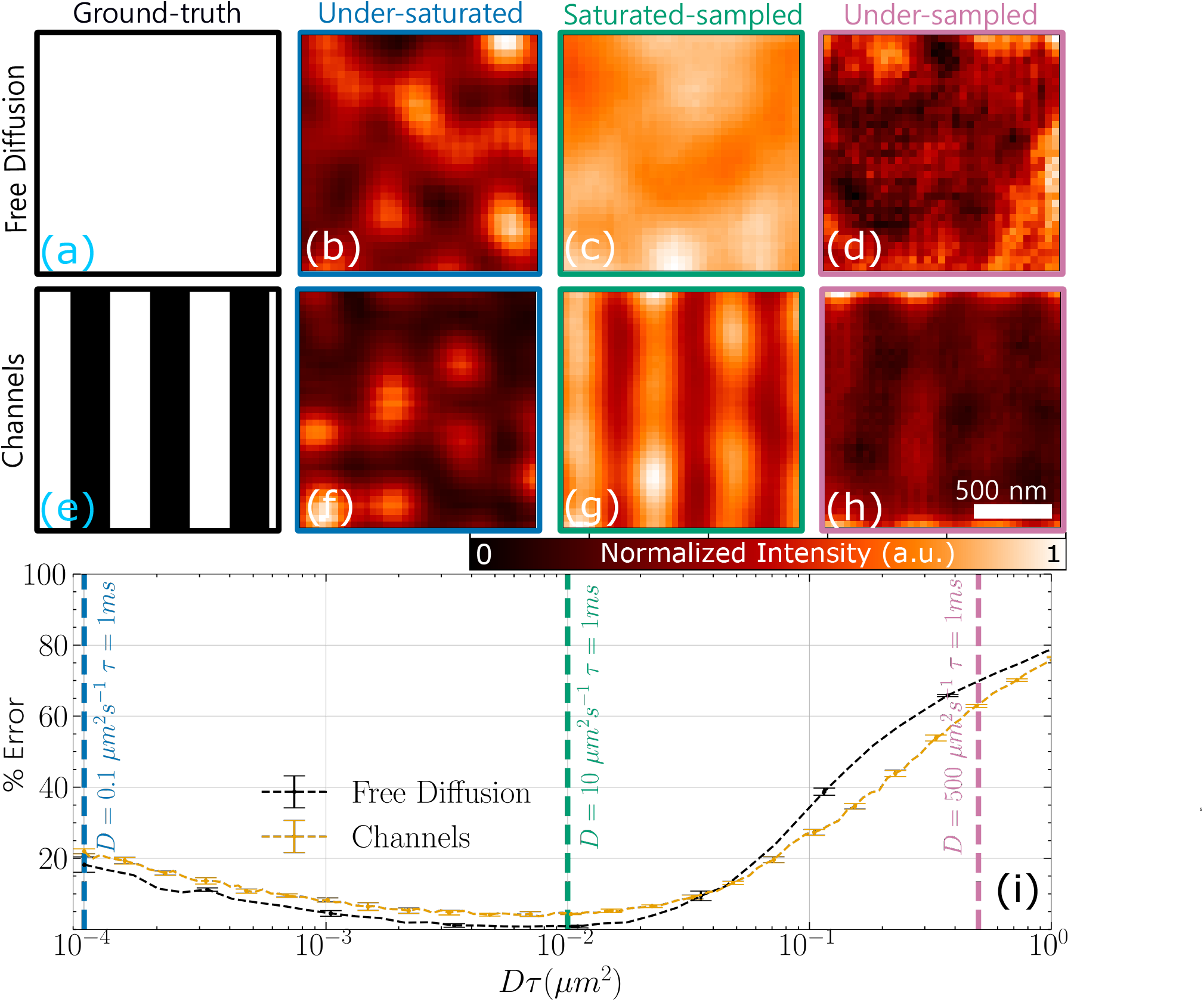
Simulating varied diffusion rates in different environments determines the range at which *XC*_2_ SOFI can applied accurately. *XC*_2_ SOFI reconstruction of 20 simulated emitters diffusing in free diffusion **(a)** and 0.304 *µm* narrow channel **(e)** structures in the under-saturated **(b, f)**, saturated-sampled **(c, g)**, and under-sampled **(d, h)** regimes. **(j)** The percent error in the calculated distance factor as a function of *Dτ* for both free diffusion and structured channels along with dashed vertical lines at the three simulated values of *Dτ* in the structures. The lowest percent error results in the most accurate SOFI images (green outlines) as emitters can fully saturate the sample and be properly sampled by the sampling rate. Each point and the associated error bar in **(i)** is calculated from the error mean and standard deviation of five separate simulations with the same parameters. The dashed vertical lines represent the three values of *Dτ* simulated in the shown structures. To achieve the SOFI images across all three regimes we simulated emitters with diffusion coefficients of 0.1, 10, and 500 *µm*^2^*/s* respectively with a exposure time (sampling rate) of 1 ms over 3000 frames. The SOFI intensity within each image is normalized to 1. Scale bar of 500 *nm*.

Our simulation and computational technique allow us to test a wide range of parameters much more efficiently than preparing hundreds of samples. However, to validate the results from the simulation, we also compare SOFI results to various experimental samples with known parameters, as detailed in our previous publications. ^23–25^

### Experimental data

Briefly, the experimental details are summarized here. A 5 *µ*L volume of a 1:5 diluted solution of stock 100 nm carboxylate-modified polystyrene beads (TetraSpek, Invitrogen) was deposited onto a nanochannel patterned coverslip (25 mm NanoSurface, Curi Bio), covered with a plain coverslip (no. 1.5 25 mm, Corning), sealed with clear nail polish (Insta-Dri, Sally Hanson, CVS), and then imaged. Imaging was performed using our home-built total internal reflection fluorescence (TIRF) microscope, as described elsewhere. ^24,33^ The video was collected at an exposure time of 5 *ms* for 60,000 frames.

## Results and Discussion

### The diffusion coefficient and exposure time determine the accuracy of the SOFI image

The values of *D* and *τ* affect the reliability and accuracy of SOFI analysis with diffusing emitters. As seen in Eq. 3, larger values of *Dτ* can reduce the resolution improvement in SOFI due to a limited reduction in the *σ* of the PSF. In addition, others have shown that blinking emitters with the addition of small amounts of diffusion introduce artifacts into SOFI images. ^29^ Understanding the effect of purely diffusing emitters on SOFI analysis is important to determine the dynamic range in which fcsSOFI can be applied.

By using Eq. 6, we find and compare the analytical and numerical distance factors to determine the accuracy in a simulated SOFI image with a known ground truth. As mentioned previously, the distance factor can be used to calculate the *σ* of the PSF. Since our distance factor in Eq. 6 has the additional term *Dτ* from diffusion as compared to stationary, blinking-based SOFI, we are no longer solving for *σ* but rather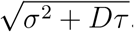 . When capturing diffusion, it is known that the right sampling rate (*τ*^−1^) must be met to capture different diffusion speeds in methods such as FCS.^26^ To find a range in which SOFI is able to use diffusion for super-resolution imaging, we compare the predicted value of 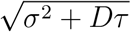 from the SOFI analysis to those from known simulated data. The percent error at different values of *Dτ* are used as a quantitative way to measure the accuracy of a SOFI image in different diffusion regimes.

From our simulations, we find that the value of *Dτ* will change the saturation quality - how well the emitters have fully diffused throughout and saturated the region with signal - in addition to the sampling quality - how well the experimental setup can sample the diffusion of emitters. When changing the value of *Dτ* there are three different regimes: the under-saturated region (Figure 3b, f), the saturated-sampled region (Figure 3c, g), and the under-sampled region (Figure 3d, h). The cut-off values of *Dτ* between these regimes vary from sample-to-sample depending on the structure simulated and a variety of other parameters. The rest of this section is devoted to describing the cause of the regimes as well as how the cut-offs between regimes change.

The under-saturated region is found in samples with slow diffusion coefficients and fast collection frame rates leading to a low value of *Dτ* . When imaging a sample with a very small value of *Dτ*, emitters are not able to diffuse throughout and fully saturate the pores in the sample. Many pores will go unvisited by emitters, leading to an under-saturated image. The error in the calculated distance factor increases, (Figure 3j), and the resulting SOFI images are spotty (Figure 3b, f), with a low number of bright areas compared to the ground truth where either the entire area (Figure 3a) or linear pores (Figure 3e) should be apparent. Exact values of *Dτ* at which SOFI analysis results in under-saturation will be dependent on the specific imaging conditions and sample (see Figure S1 in SI for examples). We find for our conditions of 3,000 frames, the regime occurs at approximately *Dτ <* 10^−3^*µm*^2^. The upper *Dτ* bound of the under-saturated region can be largely reduced by acquiring more frames and imaging over a longer period of time, allowing for emitters to diffuse throughout the sample. Collecting more frames allows for image collection at smaller values of *Dτ* while still avoiding the under-saturated region as the cut-off value of *Dτ* for the under-saturated region becomes smaller with more frames (See Figure S1). Long collection periods to accommodate fcsSOFI analysis on systems with small values of *Dτ* can be beneficial because as *Dτ* → 0 in Eq. 4, the resolution improvement approaches 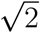 as with stationary emitters.

The under-sampled region is found in samples with fast diffusion coefficients and slow collection frame rates leading to a large value of *Dτ* . A very large value of *Dτ* leads to an experimental setup that can not fully capture the diffusion being imaged. The error calculated from the distance factor rapidly increases in the under-sampled region (Figure 3j), and we receive distorted and noisy SOFI images with very low signal (Figure 3d, h). Suppose the camera is unable to fully capture and sample the diffusion because the emitters are moving faster than they are being imaged. In that case, the fluctuations in Eq. 2, will be approximately zero, *δF* ∼ 0, causing the SOFI analysis to not return a reliable structure. The cut-off value of *Dτ* at which the under-sampled region begins varies based on the sampled structure, the pixel size of the camera, and the PSF size, but is typically between 0.01 and 0.1 *µm*^2^. Keep in mind that the numbers cited here are for conditions that we have simulated, with conditions seen in Figure S1 and S2.

The saturated-sampled region is found in samples with medium diffusion coefficients and collection frame rates leading to an intermediate value of *Dτ* (1 × 10^−3^ *< Dτ <* 2 × 10^−1^*µm*^2^ for 3,000 frames in our conditions). A value of *Dτ* between the under-saturated and under-sampled regions will lead to the most accurate SOFI image possible for a given system as the emitters have a large enough dwell time to be sampled, but small enough dwell time to diffuse throughout the structure. The emitter fluctuations are properly captured and the *XC*_2_ SOFI values yield super-resolution information. In the saturated-sampled region, the error calculated by the distance factor is near zero (Figure 3j) and the SOFI image received is clear of artifacts and resembles the ground truth (Figure 3g). The minimum error achieved in the saturated-sampled region is dictated largely by the structure being imaged. Eq. 3 and Eq. 4 both assume a constant structure to infinity, which is not always true. As the structure becomes more complex with smaller features, the approximation of integrating to infinity becomes less accurate. In Figure 3j, we can see that emitters with no structure constraints have a slightly smaller minimum distance factor error than emitters diffusing through channels (For a larger variety of sample structures, see Figure S2 in SI).

Putting all three regions together, we can see the full dynamic range of SOFI when applied to emitters which are not blinking, but rather diffusing. Due to the large variability from sample-to-sample and a large variety of experimental parameters, we cannot quote exact cut-offs for any region. However, our simulations, equations, and data provide others with some *Dτ* reference cut-offs between regions based on our trials, which can be found from the figures presented here in the SI (Figure S1 and Figure S2). Vandenberg and Dedecker’s previous work on simulated diffusing fluorescent proteins with a significant on-time ratio of 9% found no noticeable distortions for *Dτ* ≤ 0.03 *µm*^2^.^29^ We find similar results as 0.03 *µm*^2^ lies within the saturated-sampled regime in Fig. 3 for typical samples. Our work allows for a needed and more complete understanding of SOFI analysis on emitters that do not blink (on-time ratio of 100%), and our results emphasize the importance of understanding the fluctuations of your sample. A strong understanding of your experimental conditions will determine an appropriate range for *Dτ* . When imaging samples that differ largely from those presented here, it is strongly recommended to use the scripts we have provided on GitHub to find more precise cut-offs between regions. ^32^ Since SOFI is being used within the saturated-sampled region, SOFI (often applied within fcsSOFI in this situation) is a viable technique for resolving super-resolution structures with optical microscopy using diffusing emitters.

### Second order cross-correlation, *XC*_2_, reduces aliasing in super-resolution images

In super-resolution imaging, difficulty in resolving nanoscale features can arise from the limited, finite pixel-size of the wide-field detector. ^17^ For example, a spatial resolution of 100 nm could not be achieved from a 200 nm pixel size, which will be pixelated. One of the main benefits of *XC*_2_ analysis is a four-fold increase in pixel density in each dimension.^13^ As the pixel density increases, the sampling rate of the resolved structure also increases. The Nyquist sampling theorem states that if a signal has a frequency lower than half the sampling frequency, then aliasing can occur.^34–36^ Structural aliasing results in features such as jagged edges, misrepresentation of the shape, and larger features than the true physical size. In the frequency domain, larger features appear at lower frequencies and lack the higher frequency components of the true, smaller feature size. Theoretically, cross-correlation of the *n*^*th*^ order allows for *n* times smaller features to be sampled without aliasing.^13,30^ With a maximum resolution improvement of 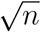 for any SOFI analysis, *XC*_2_ allows all structures uncovered by a resolution improvement to be free of aliasing, as shown in Figure 4.

**Figure 4:**
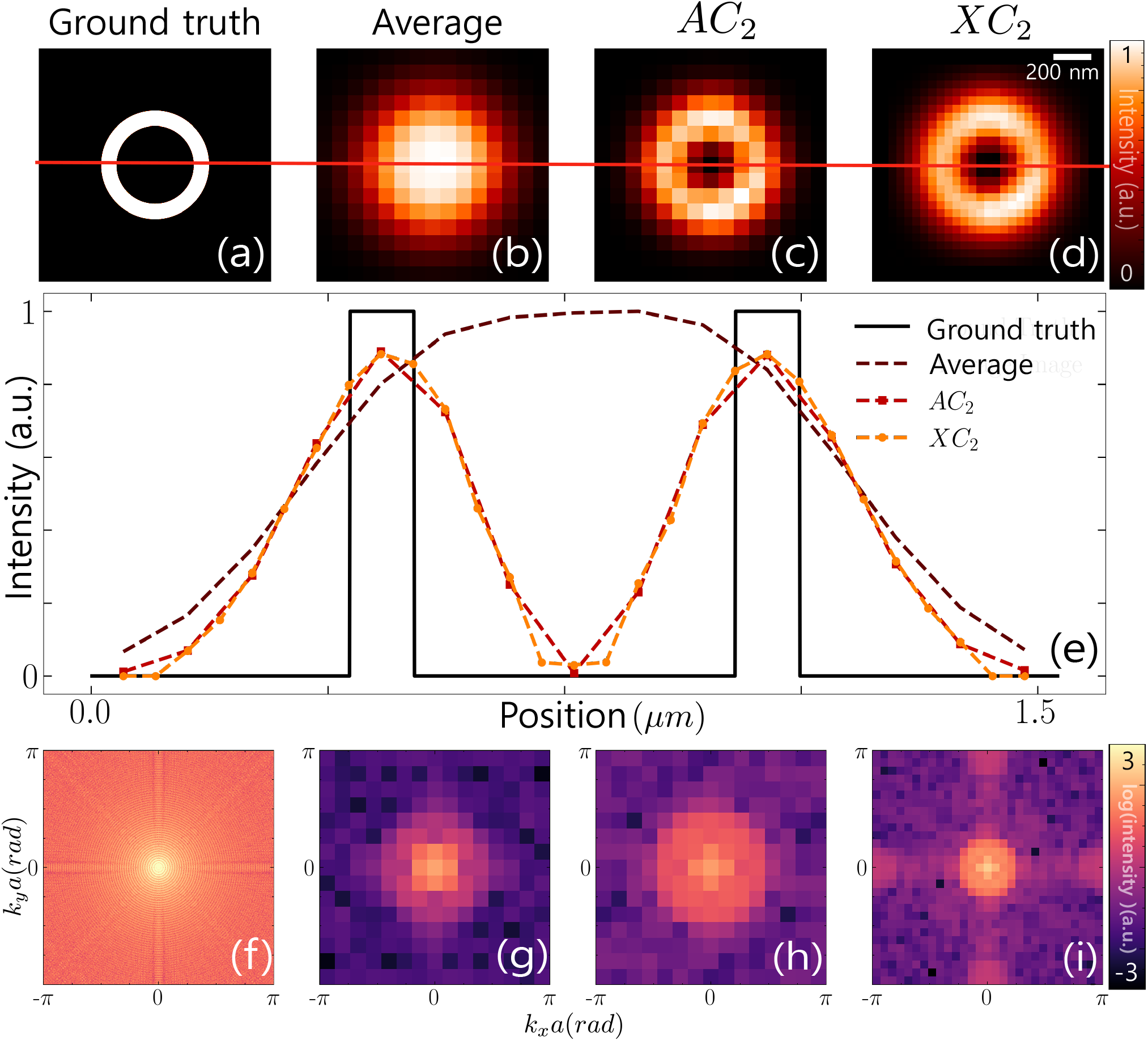
SOFI analysis in spatial and frequency domain demonstrate *XC*_2_ achieves higher sampling and recovers high frequency information. Simulated data comprised of four emitters diffusing in a ring structure at *D* = 10 *µm*^2^*s*^−1^, a time lag *τ* of 1 ms, for 30,000 frames. The size of each image is 1.22 *µm* x 1.22 *µm* and the scale bar is 200 *nm*. Spatial information from the simulated 1500 pixels x 1500 pixels binary ground truth pore structure **(a)**, 100 pixels x 100 pixels diffraction-limited average image **(b)**, 100 pixels x 100 pixels auto-correlation SOFI image (*AC*_2_) **(c)**, and 200 pixels x 200 pixels cross-correlation SOFI image (*XC*_2_) **(d)** are shown. A line section from each image in **(e)** shows a higher sampling rate in the *XC*_2_ image compared to the *AC*_2_ image. Discrete Fourier transforms are mapped for the ground truth **(f)**, diffraction-limited average image **(g)**, *AC*_2_ image **(h)**, and *XC*_2_ image **(k)**. Note that two-dimensional spatial frequency information *k*_*x*_, *k*_*y*_ is scaled by the pixel size *a* corresponding to each structure for simplicity of comparison.

Higher pixel density from *XC*_2_ analysis leads to a better representation of the true structure due to a reduction in aliasing in simulations with complex structures and a known ground truth (Figure 4). To easily see the aliasing reduction from *XC*_2_, we simulated emitters diffusing inside the ring structure in Figure 4a which yields the diffraction-limited average image in Figure 4b with no ring structure present. The *AC*_2_ SOFI image shows a ring-like structure in Figure 4c, however, the inner portion of the ring presents the structural aliasing features of sharp edges due to the low sampling. Contrasting the inner ring portion of the *XC*_2_ SOFI image in Figure 4d to the *AC*_2_ SOFI image in Figure 4c, we see a smoother, more ring like edge with less structural aliasing. The doubled sampling rate which leads to the reduction of structural aliasing in a *XC*_2_ SOFI image can be seen visually through the doubling in the number of the points in a line section plot as seen in Figure 4e. An image produced using *XC*_2_ SOFI (Figure 4d) presents a qualitatively defined and distinct image when compared to the diffraction-limited average image (Figure 4b) and the *AC*_2_ SOFI image (Figure 4c) through the reduction in structural aliasing.

Fourier maps of the SOFI images show the benefit of *XC*_2_ in resolving high-frequency information (Figure 4f-i). To briefly demonstrate the use of the Fourier map, one can look at two regions: the low-frequency region (*k*_*x*_*a, k*_*y*_*a* around (0,0)) and the high-frequency region (|*k*_*x*_*a*|, |*k*_*y*_*a*| near *π*). The low-frequency region demonstrates a constant intensity between each pixel in Figure 4a-d, where the power spectrum of SOFI signal will be highly detected in this region. The benefit of having information in the high-frequency region is to detect the edge of the structure (demonstrated where the intensity of the ring rim changes from 1 to 0). Both frequency regions are essential to construct an image. If only a high signal in the low-frequency region is present, the resulting image can be blurry. Similarly, having too much high-frequency information will result in images with high contrast. In our simulation, the average image (Figure 4g) has little high-frequency information compared to the ground truth and correlated images. We can also see that *AC*_2_ SOFI (Figure 4h) does not pronounce the cross-hair feature shown in the ground truth (Figure 4f) while *XC*_2_ SOFI does. The comparisons of the Fourier maps indicate that *XC*_2_ SOFI can recover more high-frequency information and is more comparible to the features of the ground truth. Therefore, using Fourier analysis, we can quantitatively see that *XC*_2_ SOFI helps uncover high-frequency portions of the true structures that *AC*_2_ SOFI does not resolve.

We must be careful in assessing the benefits of *XC*_2_ as one may think *XC*_2_ analysis could resolve a higher resolution structure compared to both the diffraction-limited average image and the *AC*_2_ image due to the increased sampling rate. However, the resolution in *XC*_2_ SOFI is still limited by the size of the PSF. *XC*_2_ SOFI only increases the spatial sampling rate and does not shrink the size of the PSF any more than *AC*_2_ SOFI, as shown in Eq. 6. *XC*_2_can only increase the sampling rate of structures that have already been uncovered by at most 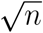 resolution improvement from any *n*-order SOFI analysis. In the line section plot of Figure 4e, we can see that although we have double the sampling, no resolution improvement is observed. The transition between high intensity and low intensity occurs at a similar rate between both the *AC*_2_ and *XC*_2_ SOFI images. *XC*_2_ SOFI reduced the aliasing through better sampling, but *XC*_2_ SOFI does not improve the resolution more than the original 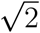 improvement from *AC*_2_ SOFI.

### Improved fcsSOFI with *XC*_2_ spatially characterizes structures in biorelevant nano-channels

To show that fcsSOFI with the *XC*_2_ approach is robust and works with real imaging conditions, here we analyze our previous experimental results of beads diffusing within nanochannels.^24^ The nano-structured channels demonstrate the effect of chemical and physical properties of the ECM in cell culture, such as fibrillar collagen, and maximize the efficiency of cell focal adhesion.^37–40^ As the analysis of the dynamics has been done using imaging FCS and cumulative distribution function (CDF), shown in Figure 1d, we only focus on the spatial resolution based on the known diffusion coefficient reported in our previous work.^23,25^ Using the fcsSOFI model as a metric shows the power to measure the diffusion rate and characterize the structure with known nanoscale channel dimensions of 800 nm x 800 nm x 600 nm (width, spacing, depth). Figure 5a and 5b demonstrates the reconstruction of the fcsSOFI image using the experimental data.

**Figure 5:**
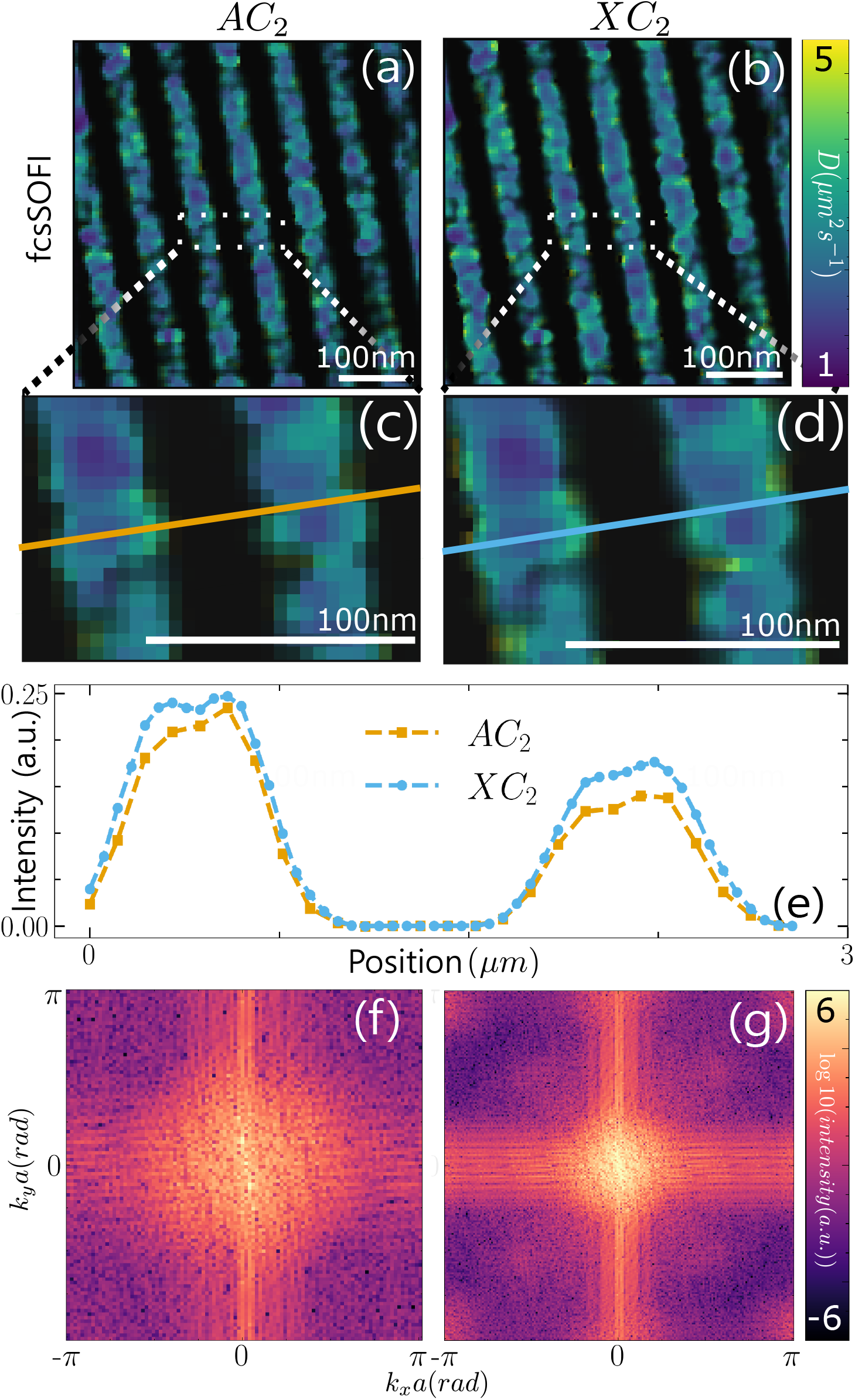
fcsSOFI analysis for diffusing beads in the nanopatterned surface using *AC*_2_ **(a)** and *XC*_2_ **(b)**. To show more features of cross-correlation analysis, we present a zoom-in image for *AC*_2_ **(c)** and *XC*_2_ **(d). (e)** Presents a line section for SOFI *AC*_2_ and *XC*_2_ image across the middle of both **(c)** and **(d)**. Then, we applied the Fourier transform to SOFI *AC*_2_ **(a)** and *XC*_2_ **(b)** information to present the benefit of higher pixel density, presented in **(f)** and **(g)** for *AC*_2_ and *XC*_2_, respectively.

*XC*_2_ SOFI analysis with dynamic information effectively increases the number of pixels while retaining high spatial frequency information under experimental conditions, providing greater detail. In this experiment, the raw wide-field image has the size of 10 *µm* x 10 *µm*, which is significantly larger than the previously shown simulated ring structure(Figure 4) in 1.5 *µm* x 1.5 *µm* region. The visual improvement of fcsSOFI with *XC*_2_ (Figure 5b) is not easily perceivable compared to that of *AC*_2_ (Figure 5a). By zooming into the fcsSOFI image, we can observe a distinction between *AC*_2_ (Figure 5c) and *XC*_2_ (Figure 5d) due to the higher pixel density. The higher sampling in the line section (Figure 5e) of *XC*_2_ presents a smoother transition between pixels, which helps reduce aliasing for samples with smaller features. Similar to the simulations, by applying a two-dimensional Fourier transform to the SOFI information, we can observe information about the high-frequency region through a spatial frequency map of both the *XC*_2_ SOFI (Figure 5g) and the *AC*_2_ fcsSOFI (Figure 5f) images. By comparing the two, we can see a recovery of the cross-hair feature inherent to high-frequency information in both images. However, the feature is stronger in the *XC*_2_ fcsSOFI image than in the *AC*_2_ fcsSOFI one. Therefore, the increased sampling of *XC*_2_ recovers the sharp, high-frequency transitions between the nanoscale channels and walls.

## Conclusion

We have shown proof that cross-correlation (*XC*_2_), analysis can be applied to diffusion-based single-molecule information. However, different experimental conditions affect a viable range of *Dτ* (speed of the molecules and camera exposure time) in which SOFI analysis can be used with dynamic emitters to fully recover the structure. We first demonstrated a regime where we could sample saturated nanostructures through the *XC*_2_ analysis at multiple diffusion coefficients and times resulting in a *Dτ* ranging from 1 × 10^−3^ − 2 × 10^−1^*µm*^2^. Next, we explored the benefit of having higher sampling due to the nature of *XC*_2_ analysis. We applied Fourier transforms to demonstrate that *XC*_2_ SOFI contains high spatial frequency values, which can be understood as the sharp change in intensity between pixels. We then demonstrated accurate results from the application of the improved fcsSOFI technique implemented with *XC*_2_ on experimental data from 100 nm beads diffusing in a nano-channel structure. In future work, we will apply our fcsSOFI technique with *XC*_2_ to understand chemical and biomedical questions, such as studying proteins in complex extracellular matrices in order to demonstrate how confinement affects diffusion dynamics.

## Supporting Information Available

The full derivation of *XC*_2_ and *AC*_2_, fcsSOFI data analysis, and the effects of structures and frames are given in the Supporting Information.

## Author Contributions

J.A. and B.W. contributed equally. Chatterjee’s current Address: The department of Biochemistry and Molecular Biology, University of Florida, Gainesville, FL, 32610

## Acknowledgement

The authors acknowledge NIH NIGMS grant R35GM142466 for financial support of this work. The authors also thank the Kisley research group, Vignesh Venkataramani, Ricardo Monge Neria, Prof. Michael Hinczewski for the helpful discussion.

