## Supporting Information for "Cross-correlation increases sampling in diffusion-based super-resolution optical fluctuation imaging"

§*ORCID: 0000-0002-8286-5264*

### Contents

1 **Derivation for $AC_2$ and $XC_2$ SOFI for diffusion-based emitters and distance factor** **S-3**

2 **fcsSOFI data analysis** **S-6**

3 **SOFI Regime Cutoff Values** **S-7**

**Figure S1** **S-7**

**Figure S2** **S-9**

**References** **S-10**

### 1 Derivation for $AC_2$ and $XC_2$ SOFI for diffusion-based emitters and distance factor

To derive the expression for both $AC_2$ and $XC_2$, we need to understand ergodicity and stationarity in the diffusion process. For the diffusion of random-walked particles, we know that the solution from the diffusion equation of particles per unit area, or concentration of fluorophores, $B$ with diffusion coefficient $D$ at position $x$ and time $t$ will yield a normal distribution function $\phi(x,t) = \frac{1}{\sqrt{4\pi Dt}} \exp -\frac{x^2}{4Dt}$. The diffusion process will meet the conditions for ergodicity and stationarity so that we can show the connection between the temporal average and the spatial average in the infinite time limit[1]

$$\langle B(x,t) \rangle_t = \frac{1}{T} \lim_{T \to \infty} \int_0^T dt B(x,t) = \langle B(x,0) \rangle \tag{S1}$$

Then, we can calculate the correlation

$$\begin{aligned}
\langle \delta B(r_1,t) \delta B(r_2, t+\tau) \rangle_t &= \langle \delta B(r_1,t) \int_{-\infty}^{\infty} dx \phi(r_2 - x, \tau) \delta B(x,t) \rangle_t \\
&= \int_{-\infty}^{\infty} dx \langle \delta B(r_1,t) \delta B(x,t) \rangle_t \phi(r_2 - x, \tau) \\
&= \int_{-\infty}^{\infty} dx \langle \delta B(r_1,0) \delta B(x,0) \rangle \phi(r_2 - x, \tau) \\
&= \int_{-\infty}^{\infty} dx \langle \delta B(r_1,0)^2 \rangle \delta(r_1 - x) \phi(r_2 - x, \tau) \\
&= \langle \delta B(r_1,0)^2 \rangle \phi(r_1 - r_2, \tau) \\
&= \langle B \rangle \phi(r_1 - r_2, \tau)
\end{aligned} \tag{S2}$$

The trick behind the relation between $\phi(x,t)$ and $\delta B(r, t+\tau)$ comes from the convolution yielded from solving diffusion equation with the boundary condition that depends on the initial fluctuation when $\tau = 0$. Also, in the original derivation of FCS,[2] the correlation of probability fluctuation will yield the last line in equation S2. The term $\langle B \rangle$ is the spatial average concentration of emitters. Furthermore, we can further rewrite the term $\langle \delta B(r_1, 0)^2 \rangle$ simpler. For this model, the concentration variance is equal to its average because we assume that the fluorescence intensity from the concentration of emitters follows a Poisson distribution.

To obtain the ratio of the $AC_2$ and $XC_2$, we need to calculate the correlation, treating that the fluorescence signal on the sCMOS comes from the contribution of all three dimensions. By symmetry, one can do the FCS calculation of the three-dimensional diffusion[1] via

$$\begin{aligned}
AC_2(r, \tau) &= \langle \delta F(r, 0) \delta F(r, \tau) \rangle_t \\
&= \int_{-\infty}^{\infty} dr_1 \, U(r - r_1) \int_{-\infty}^{\infty} dr_2 \, U(r - r_2) \epsilon_1 \epsilon_2 \left\langle \delta B(r_2, t + \tau) \delta B(r_1, t) \right\rangle_t \\
&= \int_{-\infty}^{\infty} dr_1 \, U(r - r_1) \int_{-\infty}^{\infty} dr_2 \, U(r - r_2) \epsilon_1 \epsilon_2 \frac{\langle B \rangle}{(4\pi D\tau)^{3/2}} \exp \frac{-(r_1 - r_2)^2}{4D\tau} \\
&= \epsilon_1 \epsilon_2 \frac{\langle B \rangle}{(4\pi D\tau)^{3/2}} \left( \frac{\pi}{\frac{1}{2\sigma^2} + \frac{1}{4D\tau}} \right)^{3/2} \int_{-\infty}^{\infty} dr_1 \exp \left[ \frac{-(r_1 - r)}{2\sigma^2}^2 + \frac{-(r_1 - r)^2}{2(\sigma^2 + 2D\tau)} \right] \\
&\sim \left( \frac{1}{1 + \frac{\tau}{\tau_D}} \right)^{3/2}
\end{aligned} \tag{S3}$$

where $\tau_D = \frac{\sigma^2}{D}$. To simplify our calculation, we treat the size of PSF in each dimension to be the same. In traditional FCS calculation, the PSF in the z-axis would differ from the x-y plane. By symmetry, the integrals of each dimension are independent. One could achieve the conventional three-dimensional FCS equation by changing the standard deviation of the PSF profile in the axial dimension. Also, intensity $\epsilon_1$ and $\epsilon_2$ and the average concentration $\langle B \rangle$ are independent of position and can moved out of the integration. The limitation of ignoring these parameters is that we might not be able to obtain the bulk parameter like a traditional FCS calculation. However, we can ignore this calculation to obtain the distance factor. Next, we will follow the exact derivation from $AC_2$ for $XC_2$.

$$
\begin{aligned}
XC_2(r_A, r_B, \tau) &= \langle \delta F(r_A, 0)\delta F(r_B, \tau)\rangle_t \\
&= \int_{-\infty}^{\infty} dr_1\, U(r_A - r_1) \int_{-\infty}^{\infty} dr_2\, U(r_B - r_2)\epsilon_1\epsilon_2 \left\langle \delta B(r_2, t+\tau)\delta B(r_1, t)\right\rangle_t \\
&= \int_{-\infty}^{\infty} dr_1\, U(r_A - r_1) \int_{-\infty}^{\infty} dr_2\, U(r_B - r_2)\epsilon_1\epsilon_2 \frac{\langle B\rangle}{(4\pi D\tau)^{1/2}} \exp \frac{-(r_1 - r_2)^2}{4D\tau} \\
&= \epsilon_1\epsilon_2 \frac{\langle B\rangle}{(4\pi D\tau)^{1/2}} \sqrt{\frac{\pi}{\frac{1}{2\sigma^2} + \frac{1}{4D\tau}}} \int_{-\infty}^{\infty} dr_1 \exp \left[ \frac{-(r_1 - r_A)^2}{2\sigma^2} + \frac{-(r_1 - r_B)^2}{2(\sigma^2 + 2D\tau)} \right] \\
&\sim \exp - \frac{(r_A - r_B)^2}{4(\sigma^2 + D\tau)} \int_{-\infty}^{\infty} dr_1\, \exp - \frac{((-2r_1 + r_A + r_B)\sigma^2 + 2D(-r_1 + r_A)\tau^2)^2}{4\sigma^2(\sigma^2 + D\tau)(\sigma^2 + 2D\tau)}
\end{aligned}
\tag{S4}
$$

With this bound in the infinitely large image, the analytical integration yields

$$
XC_2(r_A, r_B, \tau) \sim \exp - \frac{(r_A - r_B)^2}{4(\sigma^2 + D\tau)} \left( \frac{1}{1 + \frac{\tau}{\tau_D}} \right)^{3/2} \tag{S5}
$$

Unlike the results for static photoblinking emitters, our results yielded in Eq. S3 for $AC_2$ and Eq. S5 for $XC_2$ do not depend on position. So, the position to center the virtual pixels should not matter in this model. In theory, the further the distance between two pixels, the less correlation between each signal would be. This is also why we chose only the three closest pixels to do the correlation following the technique shown in Figure 1b and Figure 2d. The restrictions of the model present here will be discussed later to demonstrate the limitations of structure and dynamics. Lastly, we consider the signal from only one different $\tau$. Therefore, we can numerically obtain the value of $AC_2$ and $XC_2$ and obtain the numerical value of the distance factor.

### 2 fcsSOFI data analysis

Provided in Github,[3] the fcsSOFI code with $XC_2$ analysis includes example data and a user guide. Chatterjee et al.[4] and Kisley et al.[5] presented a full description of methods to apply second-order auto-correlation for SOFI and FCS analysis with technical information on how to create a fcsSOFI map presenting both diffusion and structural information. Here we will briefly discuss how to combine diffusion and structural information in the case of cross-correlation analysis.

Although $XC_2$ analysis increases the number of pixels by two-fold, imaging FCS from $AC_2$ is still limited by the original number of pixels in the raw data. To combine the structural information from $XC_2$ SOFI with the dynamics information from imaging FCS, we need to increase the number of pixels in the imaging FCS map. We can increase the number of pixels by assuming all particles within one pixel are diffusing at the same speed and splitting each imaging FCS pixel into four pixels each with with equivalent diffusion coefficients. We can then overlay the $XC_2$ SOFI as a transparency map on top of the imaging FCS map to create an fcsSOFI image as seen in Figure. 1d.

We report the diffusion dynamics in two different ways: a visual map of the diffusion coefficient at each pixel as calculated from imaging FCS and a weighted cumulative distribution of all diffusion coefficients. The visual map is used for creating the fcsSOFI image as described above. A weighted cumulative distribution is also reported to allow for a more qualitative analysis of the diffusion coefficients calculated within the sample. The cumulative distribution is weighted by the SOFI values. By weighting the diffusion values with the SOFI values, we are able to remove diffusion values that we know are not accurate due to extremely low SOFI values. Chatterjee et al.[4] showed the benefit of cumulative distribution functions to represent the dynamics as a function of the nanostructure. With both the fcsSOFI map and the weighted cumulative distribution function, we can get a full understanding of the dynamics within a sample.

### 3 SOFI Regime Cutoff Values

SOFI regime cut-off values can change for various reasons, such as the number of collected frames and structure. Here are a few figures that can be used as a reference to find acceptable collection parameters for your conditions.

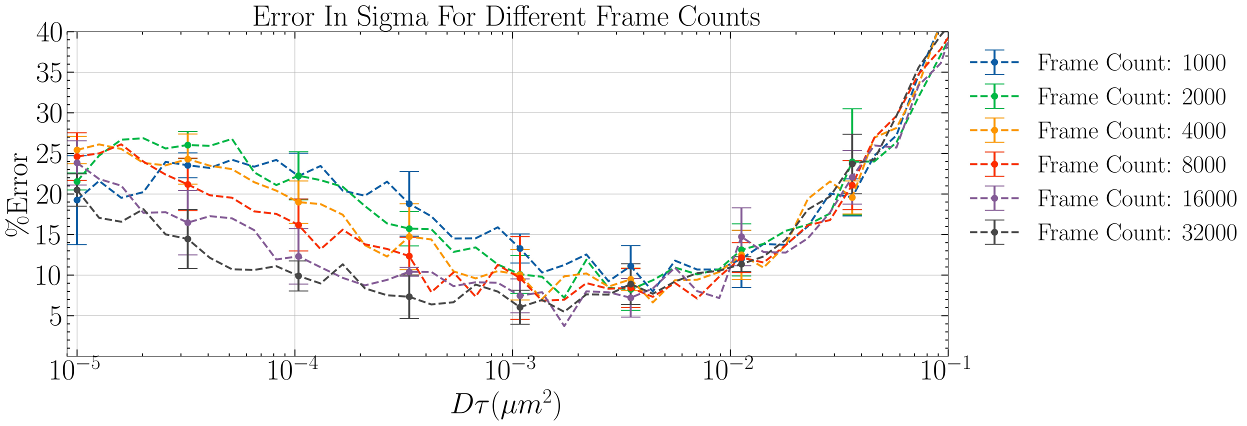


Figure S1: Plot displaying the error in the calculated distance factor as a function of $D\tau$ for multiple simulations with different frame counts. Diffusion of 15 Emitters are simulated with a PSF of 0.204 $\mu m$ and a pixel size of 0.102 $\mu m$ in a structure of randomly distributed and oriented pores with a radius mean and standard deviation of 0.510 $\mu m$ and 0.102 $\mu m$ respectively. Each point consists of the mean and standard deviation from five different simulations with the same parameters.

In Figure S1 we see the error in sigma over $D\tau$ for different frame counts. All simulations were in a structure consisting of randomly placed medium-sized pores (see Figure S1 caption for more details). As the frame count in the simulations increased from 1,000 to 40,000, the cut-off between the undersaturated regime and the saturated regime decreased. With more imaged frames, smaller diffusion, and faster frame rates can be imaged with more accuracy. With 1,000 frames, the saturated-sampled regime is within a very small range. With 40,000 frames, the saturated-sampled regime is much larger, making it easier to accurately image your sample.

In Figure S2, we see the error in sigma over $D\tau$ for different structures including channels, rings, small, medium, and large pores, and free diffusion (no structure). See the figure caption for more details about the simulation. As the structure changes, the lowest error in the calculated distance factor also changes within the saturated-sampled regime. As structures get smaller (to sizes smaller than the PSF) the minimum error achievable using SOFI with diffusing emitters increases. An increased error with smaller structures is expected since all theories were developed assuming infinite integration over a uniform structure in all spaces. As the structure gets smaller and smaller, this approximation becomes less and less accurate. When imaging smaller structures, slow diffusion coefficients and faster frame rates must be used to acquire the most accurate SOFI image.

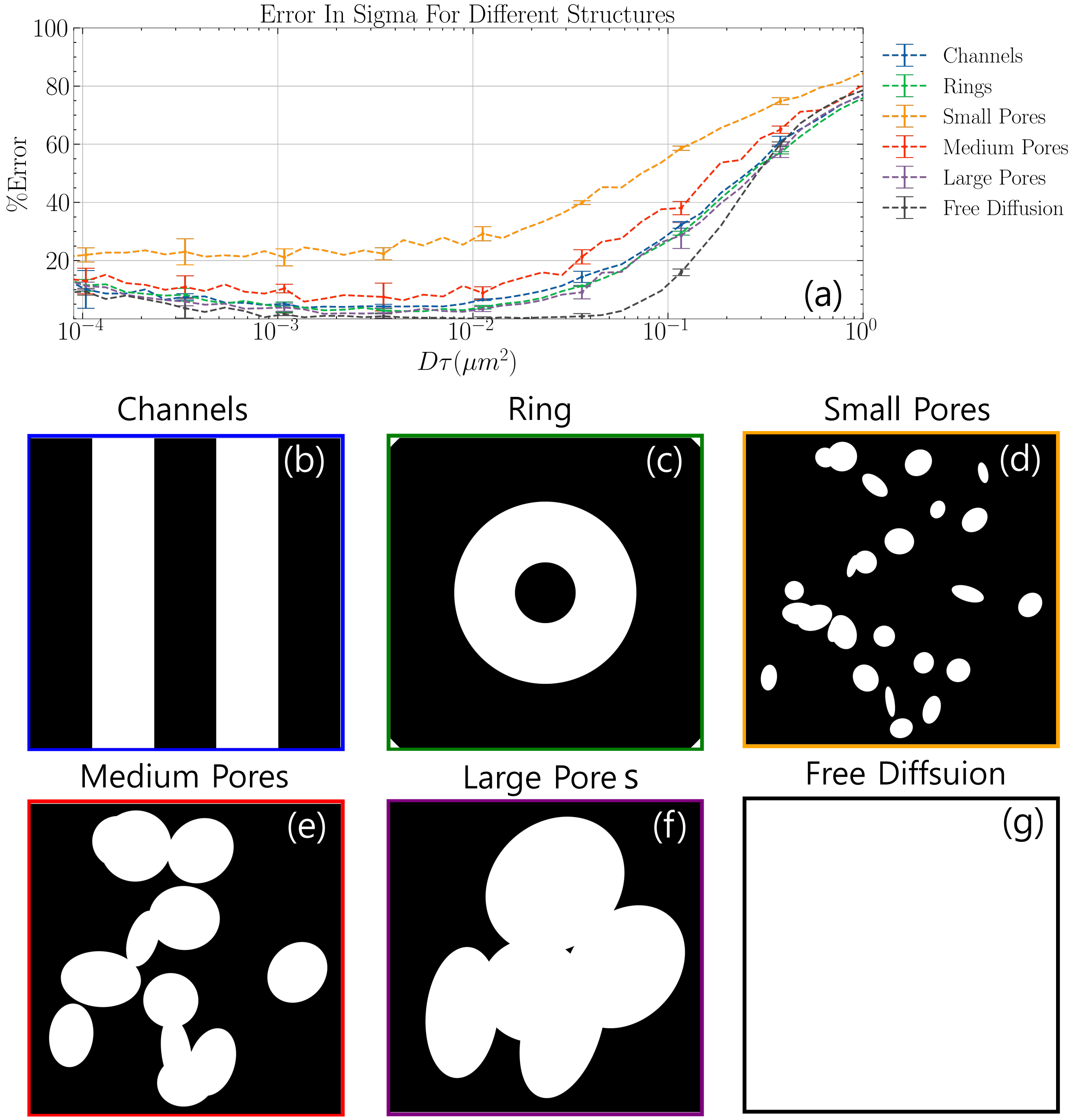


Figure S2: **(a)** displaying the error in the calculated distance factor as a function of $D\tau$ for multiple simulations with different structures (**(b-g)**. Emitters simulated with a PSF of 0.204 $\mu m$ and a pixel size of 0.102 $\mu m$. The Channels structure **(b)** contains vertical channels with a width of 1.02$\mu m$ and spaced apart by 1.02$\mu m$. The Rings structure **(c)** contains rings (or donuts) centered at the middle of the structure with a thickness and spacing of 1.02$\mu m$. The Small**(d)**, Medium**(e)**, and Large Pores**(f)** structures are made up of randomly distributed and oriented pores with a radius mean and standard deviation of 0.204 $\mu m$ and 0.051 $\mu m$, 0.510 $\mu m$ and 0.102 $\mu m$, and 1.02 $\mu m$ and 0.306 $\mu m$ respectively. The Free Diffusion structure (g) contains no structure and emitters are allowed to diffuse completely freely. Each point consists is the mean and standard deviation from five different simulations with the same parameters.
